# Changes in the wing spot pattern of the endangered butterfly *Phengaris teleius* thirty years after its reintroduction

**DOI:** 10.64898/2026.11.25.768883

**Authors:** Daniel Sánchez-García, Luca Pietro Casacci, Irma Wynhoff, Violette Chiara, Magdalena Witek

## Abstract

1. Reintroduced insect populations founded by few individuals can change in phenotype through founder effects, genetic drift and new environmental conditions. Butterfly wing spot patterns, their fluctuating asymmetry and their variability can reflect such changes and population stress.

2. We studied the spot pattern of the hindwing underside of 267 *Phengaris teleius* butterflies from the Polish source metapopulation at the time of their reintroduction to the Netherlands in 1990, and from the current Polish and Dutch metapopulations (2019–2020). We analysed the presence, size, asymmetry and fusion of the spots, and the shape and variability of the spot pattern.

3. The reintroduced Dutch metapopulation differed in spot pattern shape from both Polish ones and was less variable than the source metapopulation. The two Polish metapopulations did not differ from each other in shape. Dutch butterflies also had a more frequent spot 11 than the current Polish metapopulation and a larger spot 1 than the source ones, and the concordance between both wings in the presence of the facultative spots differed among metapopulations, being lowest in the Dutch one. Only the presence of spot 10 in females and spot fusion in males changed in the Polish metapopulation between 1990 and 2019.

4. These results agree with the founder effect and the loss of genetic variability after the reintroduction, although local adaptation or plasticity cannot be ruled out. Although based on a single reintroduction, these results suggest that spot pattern variability, measured from photographs, could complement the genetic monitoring of reintroduced insect populations.

## Introduction

Research on wing pattern and wing morphology in butterflies has been widely applied to explain ecological and evolutionary patterns (Parchem *et al*., 2007). The Lycaenidae family is one of the most species-rich groups of butterflies (Robbins, 1982), but the significance of wing colour and wing pattern is much less studied than in other butterfly families. Color and spot pattern of wings can have important functions like predation avoidance (Wourms & Wasserman, 1985), mate recognition (Rutowski & Rajyaguru, 2013) and thermoregulation (Taylor-Cox *et al*., 2020), although most of this evidence comes from other butterfly families. It has been demonstrated that in Lycaenidae, hindwing spots can play an important role as species and partner recognition signals. For instance, males of *Lycaeides idas* courted paper models of *Lycaeides melissa* females whose ventral hindwing spots had been reduced to look like those of *L. idas* as often as models of their own species (Fordyce *et al*., 2002). However, the function of the black spots of the hindwing underside of *Phengaris* butterflies is unknown. The wing pattern of lycaenids has a genetic basis (Lucas *et al*., 2018), but it can also change in response to temperature during development, as did the ventral spots of *Zizeeria maha* at the margins of its range expansion (Otaki *et al*., 2010). In *Phengaris arion* (Lycaenidae), variation in the wing melanization level and in the number of black spots on wings has been found among different populations both in Finland and in Poland (Sielezniew & Dziekańska, 2011; Väisänen *et al*., 1994), probably due to different climatic conditions.

Studying the wing spot pattern can also provide important knowledge about a population status through comparing fluctuating asymmetry (FA), the random deviations from perfect bilateral symmetry, and trait variability (i.e., wing spot pattern variability) among various populations. Indeed, FA and general trait variability can be sensitive to environmental and genetic stress (Parsons, 1992), although not always. In the butterfly *Parnassius apollo*, asymmetry did not rise and morphological variability did not decline after several bottlenecks (Habel *et al*., 2012).

*Phengaris* butterflies are obligate social parasites of *Myrmica* ants and they require two different resources for larval development; namely, specific food-plant species and *Myrmica* ant hosts (Thomas, 1980). Such complex adaptations pertaining to closely interacting species make them much more vulnerable to environmental changes. Nowadays, many butterfly populations are declining (van Swaay *et al*., 2011) and myrmecophilous butterflies including *Phengaris* spp. are those that suffer greatly (Settele & Kühn, 2009). In response to environmental changes and local population extinctions, actions including the translocation or reintroduction of threatened species to new or former sites are performed. *Phengaris* butterflies, despite their complicated life cycle, were successfully reintroduced in two cases: *P. arion* from Sweden to the United Kingdom (Thomas *et al*., 2009) and *P. teleius* from Poland to the Netherlands in 1990 (Wynhoff, 1998). The latter reintroduction was performed by introducing 86 butterflies from the metapopulation system of the Kraków region (Wynhoff, 1998).

Reintroduced populations are founded by a limited number of individuals, so they are expected to lose genetic variation through founder effects, bottlenecks and genetic drift (Nei *et al*., 1975). Genetic drift can change the mean of a morphological trait and reduce its variability in a few generations (Lande, 1976), while the new environment can select for different trait values or induce plastic responses (Stockwell *et al*., 2003). Both processes can act at the same time (Kolbe *et al*., 2012). Although 86 founders is not a small number, the reintroduced Dutch metapopulation of *P. teleius* has half of the alleles of the current Polish metapopulation, shows signs of a genetic bottleneck and has a smaller estimated effective population size (Sánchez-García *et al*., 2024). Moreover, the climate warmed in both regions after the reintroduction, and the current Polish butterflies have bigger hindwings than those from the source metapopulation in 1990 and from the Netherlands (Sánchez-García *et al*., 2024). However, the wing spot pattern of these butterflies has not been studied yet.

The thirty butterfly generations since the reintroduction give the opportunity to study possible changes and adaptations that could have occurred. As the reintroduction process is connected with changes in many habitat parameters and very often leads to a decrease in genetic variation in reintroduced populations, studying its potential effect on the butterfly wing pattern could be enlightening. The aim of our study was to test whether and how the number of black spots, their size, symmetry and fusion, and the shape and variability of the spot pattern have changed between the source and reintroduced metapopulations of *P. teleius* thirty years after the reintroduction. As we also possess the wings of butterflies from the Polish source metapopulation from 1990 (the year of reintroduction), we analysed how the wing spot patterns diverged through time between the original source metapopulation and the current reintroduced and source ones. This design allows us to compare the predictions of different processes. (1) If the changes were caused by the founder effect and genetic drift after the reintroduction, the Dutch metapopulation should differ from both the source metapopulation in 1990 and the current Polish one, these two should not differ from each other, and the spot pattern variability of the Dutch metapopulation should be lower; its FA could also be higher, if the loss of genetic variation reduced its developmental stability (Parsons, 1992). (2) Environmental changes shared by both regions, such as climate warming, would make both current metapopulations differ from the butterflies of 1990, whereas local changes in the source region would affect only the current Polish one. (3) Adaptation or plastic responses to the local conditions of the Netherlands would also make the Dutch metapopulation differ from both Polish ones.

## Material and methods

### Study site of the source metapopulation

The studied *Phengaris* (=*Maculinea*) *teleius* butterfly metapopulation occurs in the outskirts of Kraków city in South Poland (50°01’N, 19°54’E). The landscape of this area is composed mostly of abandoned or rarely managed grasslands, arable fields, forests, and settlements (Kajzer-Bonk *et al*., 2016). The habitats of the focal butterfly species are a part of a large meadow complex with an area exceeding 200 ha and consisting of several dozen nutrient-poor to mesotrophic meadows with varying densities of *Sanguisorba officinalis*, the only foodplant of caterpillars of *P. teleius*. The three investigated meadow patches, where butterflies were collected, are characterized by relatively large areas (2.4, 6.2 and 33.3 ha, respectively). Currently, the whole meadow complex faces increasing urbanization pressure, due to growing human settlement and infrastructure (Kajzer-Bonk & Nowicki, 2022). Local populations range from several hundred to a few thousand individuals (Nowicki *et al*., 2007), but since the reintroduction, habitat patches have been lost to built-up areas, afforestation and meadow succession, and the connectivity of the metapopulation has decreased (Sánchez-García *et al*., 2024).

### Study site of the reintroduced metapopulation

The nature reserve Moerputten (115 ha) is located south of the city of ’s-Hertogenbosch (The Netherlands) and covers the central part of the Natura 2000 nature reserve “Vlijmens Ven, Moerputten en Bossche Broek” (931 ha; 51°41’N, 5°15’E). On the outer borders of Moerputtennature reserve, partially within the forest, different types of grasslands are found, of which the hay meadows with a high abundance of *S. officinalis* are the most important habitat of *P. teleius*. The historical metapopulation of *P. teleius* went extinct in 1976, and *P. teleius* specimens were reintroduced in 1990 on the moist meadows (Wynhoff, 1998), at that time the only suitable site of this butterfly species in the Netherlands. The reintroduced metapopulation consisted of 33 males and 53 females of *P. teleius*. Nowadays, *P. teleius* forms a small metapopulation, isolated from any other population of the species, with one core population on the meadows at the southern border of the core reserve and two to three small populations on other meadows within the nature reserve. No further butterflies have been introduced since 1990. The Dutch metapopulation currently consists of a few thousand butterflies, and the connectivity of its habitat patches, already higher than that of the Polish metapopulation at the time of the reintroduction, sharply increased in the mid 2010s thanks to the restoration of about 250 ha of fen meadows in neighbouring reserves (Sánchez-García *et al*., 2024; Sevilleja *et al*., 2022). Mean temperature increased in both regions between 1990 and 2020, but it was lower in the Netherlands, where it currently equals that of Poland in 2000 (Sánchez-García *et al*., 2024).

### Experimental design and sampling

Data were collected from two metapopulations (Polish and Dutch) and at different times: in 1990 (the year of reintroduction) from the source metapopulation from Poland (=PL1990), in 2019 from the current metapopulation from Poland (=PL2019) and in 2020 from the reintroduced metapopulation in the Netherlands (=NL2020). Wings of individuals from PL1990 used for the analysis came from butterflies that did not survive the trip from Poland to the Netherlands for the reintroduction. They were dried and preserved (n = 63) in an entomological box. The hindwing undersides of these individuals were digitally photographed and used for morphological and spot pattern analysis (see below).

Butterflies were collected in 2019 in three closely located meadows in the Kraków region, which are the areas where in 1990 *P. teleius* adults were collected for reintroduction. In 2020, butterflies were also collected on three closely located meadows in Moerputten nature reserve. Butterflies were captured with entomological nets. After capture, each butterfly was put into a small jar and treated with carbon dioxide for ten seconds to anaesthetize it. Then the butterfly was laid on millimetric graph paper and photographs were taken from the left and right sides. The underside of both hindwings, where the studied black spots are located, was photographed using a Nikon D7200 camera and a Laowa 100 mm macro lens. Finally, the butterflies were marked with a fine-tipped waterproof Stabilo pen on the ventral part of the right forewing to prevent re-sampling of the same individual. All butterflies were released at the place of capture when they were fully awake again. The same butterflies and photographs were used by Sánchez-García *et al*. (2024) to study the hindwing size and shape of these metapopulations and the genetic structure of the current ones; their wing spot pattern, analysed here, was not studied there.

### Geometric morphometric approach

A total of 267 butterflies (PL1990 n = 63, PL2019 n = 112, NL2020 n = 92) were used for studying the wing spot pattern morphology of the hindwing underside. The spot pattern refers to the motif created by the combination of the different spots found on the wing. Up to eleven landmarks, one at the centre of each spot, were digitized in every picture with the software tpsDig v.2.32 (Bookstein, 1997; Rohlf, 2018). We considered as landmarks the points that could be precisely identified (i.e. spot location; Fig. 1; Supporting Information, Fig. S1). The position of the absent spots, mostly the facultative spots 10 and 11, was estimated with a thin-plate spline interpolation by using the estimate.missing() function of the R package geomorph (Adams *et al*., 2026; Baken *et al*., 2021); to check that the shape results did not depend on these estimated positions, the Procrustes ANOVA and the pairwise comparisons among metapopulations were repeated with the nine permanent spots only. The landmarks were used to estimate both wing spot pattern shape and centroid size, calculated as the square root of the sum of squared distances of all the landmarks from their centroid, which is the most appropriate measure of overall size (Bookstein, 1997). Additionally, thirty-one landmarks were digitized on the wing outline to calculate the wing centroid size (missing outline landmarks, in 30 wings, were estimated in the same way), which was used to test whether the spot pattern centroid size is a good estimator of wing size (see below).

**Fig. 1:**
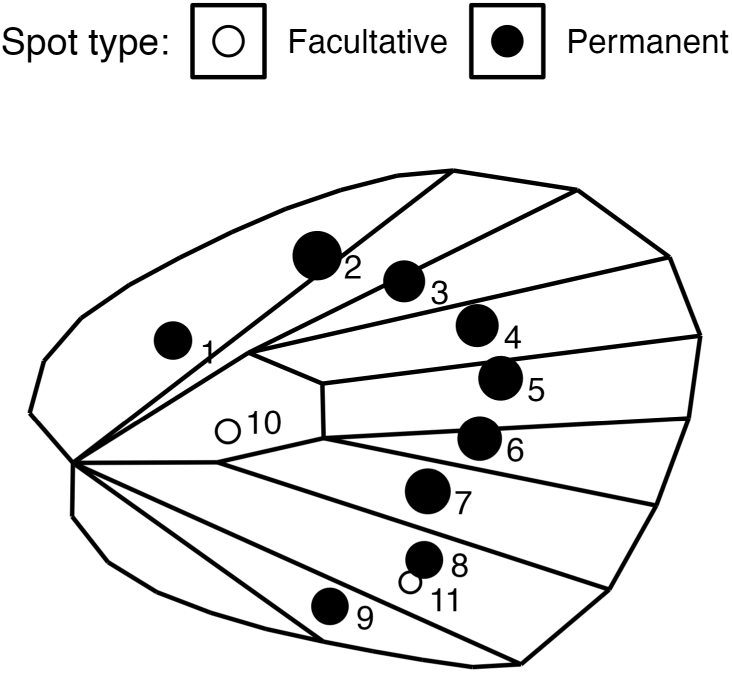
Spot pattern landmark locations in the general consensus hindwing underside of *P. teleius*. Circle area is proportional to the mean area of each spot (left hindwings of all butterflies in which it is present) and the numbers indicate the spot identification code. Filled circles: permanent spots (1–9); open circles: facultative spots (10 and 11).

Landmark oversampling for centroid size and shape analysis was tested by performing the Landmark Sampling Evaluation Curve (LaSEC) with 99 iterations by using the lasec() function of the R package LaMBDA (Watanabe, 2017). The minimum number of landmarks to reach a fit ≥ 0.95 was 4 for the centroid size and 10 for shape, so the eleven landmarks sampled the spot pattern sufficiently for both analyses.

The raw data were subsetted and only the data from the left wing were used in the analysis to avoid bias based on directional asymmetry between left and right wings. A bilateral symmetry analysis of the analysed butterflies with both hindwings digitized (127 females and 133 males) with the bilat.symmetry() function (Adams *et al*., 2026; Baken *et al*., 2021) detected it in the spot pattern shape of both sexes (females: P = 0.025; males: P = 0.008) but not in its centroid size (females: P = 0.733; males: P = 0.967). To obtain shape variables from landmark data, a single Generalized Procrustes Analysis (GPA) of both sexes was performed using the gpagen() function (Adams *et al*., 2026; Baken *et al*., 2021); after the outlier was removed, the aligned coordinates were split by sex. Outliers were identified with the plotOutliers() function, which orders the individuals by their Procrustes distances from the mean shape (Adams *et al*., 2026; Baken *et al*., 2021). Specimens whose Procrustes distance exceeded the upper quartile plus 1.5 times the interquartile range of the (Box-Cox transformed) distances were considered outliers and consequently removed from the dataset (n = 1, a male from PL2019).

### Computer assisted spot detection

Spot areas were measured from the hindwing images with a program developed in Python (3.9), which detected the black spots automatically and allowed the errors of the detection to be corrected by hand in a graphical user interface (Appendix S1); all Python scripts are available to download from a GitHub public repository (https://github.com/VioletteChiara/Wing_spots). The images whose automatic detection errors could not be corrected by hand (n = 12, 3 of them left hindwings) were left out of the analyses of spot presence, area and fusion, and the butterflies with an uncorrected left hindwing from all analyses; the landmark positions of the right hindwings, digitized by hand, were kept. Area was calculated in pixels and then converted to mm^2^. One spot area (spot 6 of the left hindwing of a PL2019 male) was clearly mismeasured, more than four times its area in the right hindwing although both spots looked alike in the image, and could not be measured again. It was therefore set as missing, and this individual was left out of the analyses of the total spot area and of the whole spot pattern (NMDS, PERMANOVA and dispersion).

### Morphometric statistical analysis

All statistical analyses were performed in R version 4.5.0 (R Core Team, 2025). The spot pattern centroid size was tested for correlation with the wing centroid size with a Spearman’s rank correlation by applying the cor.test() function (R Core Team, 2025) (Supporting Information, Fig. S2). After testing for centroid size correlation the data frame was subsetted and all the analyses were separately performed for females and males, since sex affected the spot pattern shape (Procrustes ANOVA with both sexes, comparing nested models with log centroid size and metapopulation as covariates, Z = 4.96, P = 0.001; the sex × metapopulation interaction was not significant, P = 0.671).

The differences in the hindwing spot pattern shape among metapopulations and allometry were tested by using the procD.lm() function (Adams *et al*., 2026; Baken *et al*., 2021). It performs a Procrustes ANOVA with permutation; we used residual randomization (RRPP) with type II sums of squares, so that the effect of size was tested after that of metapopulation and vice versa. The model was built using as predictor variables the logarithm of the spot pattern centroid size and metapopulation. Their interaction, tested by comparing nested models, was not significant (females: P = 0.192; males: P = 0.653), so a common allometric slope was assumed for all metapopulations. A pairwise comparison was also performed between metapopulations by applying an estimated marginal means (EMMs) test by using the function pairwise() of the R package RRPP (Collyer & Adams, 2026; Collyer & Adams, 2018), with Bonferroni-adjusted P values (over the three pairwise comparisons among metapopulations). The effect of allometry was removed from the pairwise comparison by using shape ∼ log(centroid size) as the null model. The spot pattern shape differences among metapopulations were represented with a canonical variate analysis (CVA) of the Procrustes shape coordinates by using the CVA() function of the R package Morpho (Schlager, 2017), and the first canonical variate was plotted against the logarithm of the centroid size to show the allometric effect (Supporting Information, Fig. S3). All shape tests used 999 permutations; the nested-model tests and the bilateral symmetry analysis randomized the raw values.

Spots 1 to 9 were present in nearly all individuals (see Results), so spot presence was only modelled for the two facultative spots (10 and 11), with a binomial generalized linear mixed model (GLMM) including metapopulation, spot identity and their interaction as fixed effects and the individual as a random intercept, since each butterfly contributes both spots, by using the glmer() function of the R package lme4 (Bates *et al*., 2026). Spot area was log-transformed and analysed with a linear mixed model (LMM) with the same fixed and random effects, including all the spots present, and with the logarithm of the spot pattern centroid size as a covariate, since spot area scaled with size; a separate residual variance was estimated for each spot, by using the lme() function of the R package nlme (Pinheiro *et al*., 2026). The total surface of the melanized black area was calculated as the sum of the area of all spots. It was log-transformed and analysed with a Gaussian generalized linear model using the metapopulation as a predictor variable and the logarithm of the spot pattern centroid size as a covariate, by using the glm() function (R Core Team, 2025). For the figures and the Bray-Curtis distances, spot areas (Supporting Information, Fig. S4) and the total spot area were standardized to the mean centroid size with the allometric slope estimated by their respective models. The spot fusion (between spots 8 and 11) was analysed only in the individuals with both spots present, with a generalized linear model with binomial distribution using the metapopulation as a predictor variable (fusion ∼ metapopulation) by using the glm() function (R Core Team, 2025).

Fluctuating asymmetry was tested for spot presence. Both hindwings (left and right) were considered for this analysis. An individual was considered asymmetrical for a spot when the spot was present in one of its wings and absent in the other one. As the proportion of asymmetrical individuals depends on how often a spot is present, asymmetry was measured as the concordance between the two wings, which does not depend on the frequency of the spot. The concordance was estimated as the odds ratio of the presence of a spot on the right wing given its presence on the left one, for each metapopulation and averaged over spots 10 and 11. A high odds ratio indicates that both wings agree, and an odds ratio of one that the presence of a spot on one wing does not predict its presence on the other. As for presence, asymmetry was only modelled for the facultative spots 10 and 11, with a binomial GLMM of the presence of each spot on the right wing including its presence on the left wing, metapopulation, spot identity and their two-way interactions as fixed effects and the individual as a random intercept, by using the glmer() function (Bates *et al*., 2026); the interaction between the presence on the left wing and the metapopulation tests whether the concordance between wings differs among metapopulations. Unlike landmark-based measures of fluctuating asymmetry, which require replicated measurements to separate them from measurement error, the presence or absence of a spot can be scored with negligible error.

The wing spot pattern distance between individuals based on spot presence, size-standardized spot area (divided by the mean spot pattern centroid size, so that it is on a scale comparable to that of presence and fusion) and fusion (between spots 8 and 11) was calculated by using the vegdist() function of the R package vegan (Oksanen *et al*., 2026) with the Bray-Curtis dissimilarity index. A permutational analysis of variance (PERMANOVA; 999 permutations) was applied to assess the significance of the metapopulation by the adonis2() function (Oksanen *et al*., 2026), and the distances were represented with a non-metric multidimensional scaling (NMDS; two dimensions) by using the metaMDS() function (Oksanen *et al*., 2026) (Supporting Information, Fig. S5). The spot pattern variability of each metapopulation was assessed as the Bray-Curtis distance of each individual to the centroid of its metapopulation, and compared among metapopulations with a permutation test of multivariate dispersion (PERMDISP; 999 permutations) by using the betadisper() and permutest() functions (Oksanen *et al*., 2026), with Bonferroni-adjusted pairwise permutation tests.

The significance of the terms of the generalized linear models was tested with type II likelihood-ratio tests by using the Anova() function of the R package car (Fox *et al*., 2026), and that of the mixed models with likelihood-ratio tests between nested models, testing the main effects in the model without the interaction. The metapopulations were pairwise-compared with estimated marginal means (EMMs) by using the emmeans() function of the R package emmeans (Lenth & Piaskowski, 2026), with Bonferroni-adjusted P values (over the three comparisons among metapopulations, within each spot in the models with spot identity), and groups were labelled in the figures with compact letter displays of these comparisons by using the R package mult-compView (Graves *et al*., 2026). Model assumptions were checked for every model (Table S1), with simulated residuals for the binomial models (R package DHARMa) (Hartig, 2026), with the distribution and variance of the residuals for the Gaussian ones, also testing total spot area with heteroscedasticity-robust (HC3) standard errors (R package sandwich) (Zeileis & Lumley, 2024), and with the homogeneity of shape disparity among metapopulations for the Procrustes ANOVAs (morphol.disparity() function (Adams *et al*., 2026; Baken *et al*., 2021)); the spot area model was refitted without its observations with extreme residuals (|normalised residual| > 3).

## Results

### Morphometric analyses

#### Correlation between wing and spot pattern centroid size

Spot pattern centroid size and wing centroid size showed a strong, significant correlation (Spearman’s *ρ* = 0.94, P < 0.001; Fig. S2), suggesting that either can be used as an estimator of wing size.

#### Wing spot pattern shape

The analysis of the wing spot pattern shape showed differences among the individuals from the source (PL1990), current Polish (PL2019) and reintroduced Dutch (NL2020) metapopulations for both females and males (Procrustes ANOVA, females: d.f. = 2, Z = 3.58, P = 0.001; males: d.f. = 2, Z = 3.52, P = 0.001; Fig. 2). Spot pattern shape also changed with centroid size (allometry), significantly in males and at the threshold of significance in females (Procrustes ANOVA, females: d.f. = 1, Z = 1.60, P = 0.050; males: d.f. = 1, Z = 2.00, P = 0.023; Fig. S3). In both sexes, the reintroduced Dutch metapopulation (NL2020) differed in spot pattern shape from both Polish metapopulations, whereas the source (PL1990) and the current Polish metapopulation (PL2019) did not differ (females: Z = 1.39, P = 0.249; males: Z = 1.93, P = 0.084; Table S2). The positions of the facultative spots 10 and 11, absent in 75 and 66 of the 266 wings, respectively, were estimated for these analyses; with the nine permanent spots only (absent, and therefore estimated, in only 7 cases), the metapopulation effect remained (females: Z = 3.22, P = 0.001; males: Z = 3.15, P = 0.001), and NL2020 still differed from both Polish metapopulations in females (against PL1990: P = 0.006; against PL2019: P = 0.015) and from PL2019 in males (P = 0.006), but no longer from PL1990 in males (P = 0.105).

**Fig. 2:**
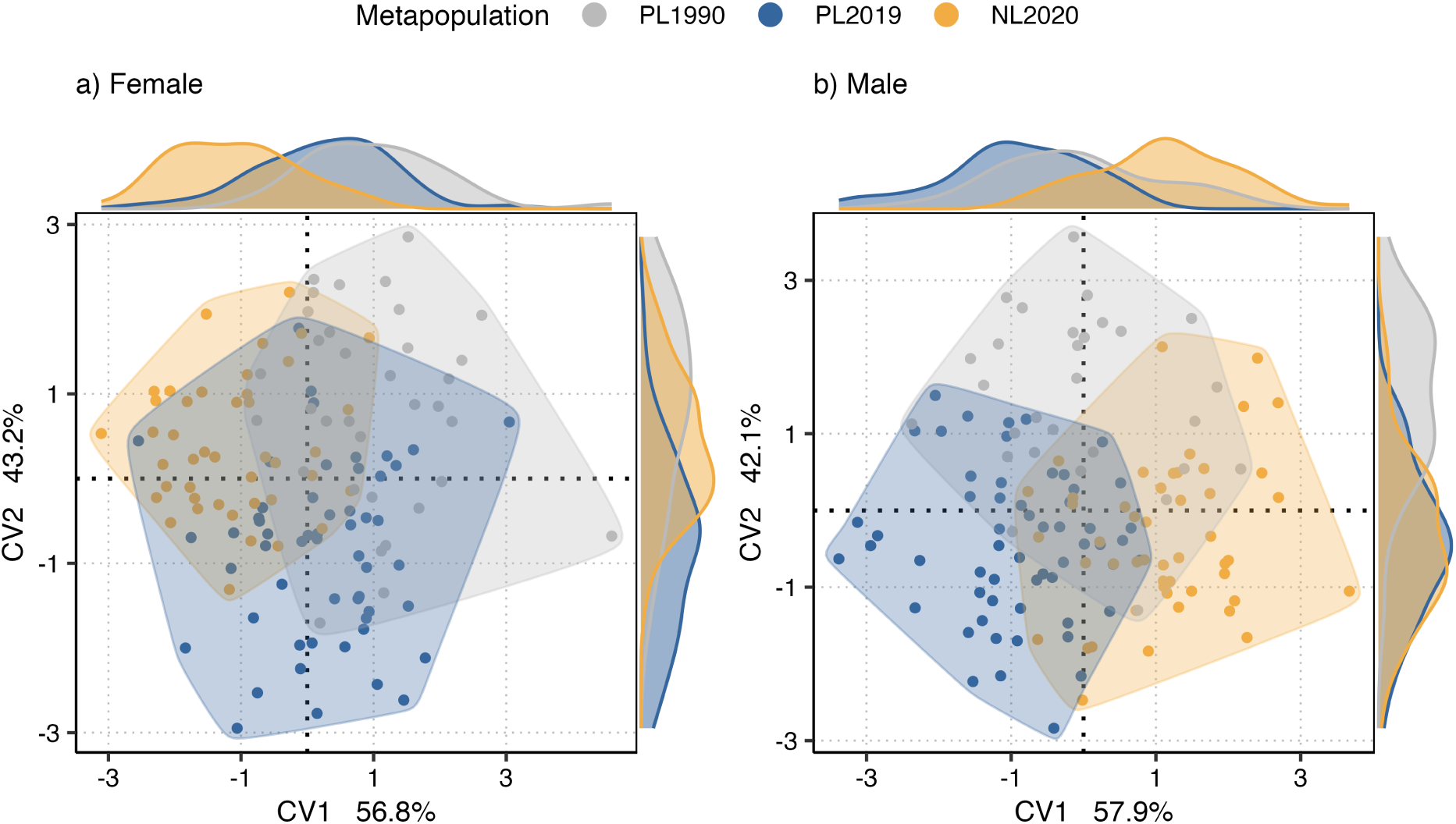
Procrustes CVA shape representation of *P. teleius* a) female and b) male wing spot pattern from the Polish source (PL1990), current Polish (PL2019) and reintroduced Dutch (NL2020) metapopulations. The axis density plots represent the distribution of each canonical variate, and the shaded areas enclose the individuals of each metapopulation.

#### Spot pattern analysis

Spots 1 to 9 were present in nearly all butterflies (at least 96% of the females and 96% of the males of every metapopulation; Fig. 3), so we consider the highly variable spots 10 and 11 as facultative spots. In both sexes, the effect of the metapopulation on the presence of the facultative spots depended on the spot (interaction, females: d.f. = 2, *χ*^2^ = 19.17, P < 0.001; males: d.f. = 2, *χ*^2^ = 19.23, P < 0.001), whereas neither the metapopulation (females: d.f. = 2, *χ*^2^ = 4.92, P = 0.086; males: d.f. = 2, *χ*^2^ = 0.31, P = 0.858) nor the spot identity (females: d.f. = 1, *χ*^2^ = 0.00, P = 1.000; males: d.f. = 1, *χ*^2^ = 1.99, P = 0.158) had a significant effect on its own. Spot 11 was significantly more frequent in the Dutch metapopulation (NL2020) than in the current Polish metapopulation (PL2019) in both sexes, and spot 10 was significantly less frequent in females from the source metapopulation (PL1990) than in females from PL2019, whereas Dutch females did not differ significantly from either (Fig. 3).

**Fig. 3:**
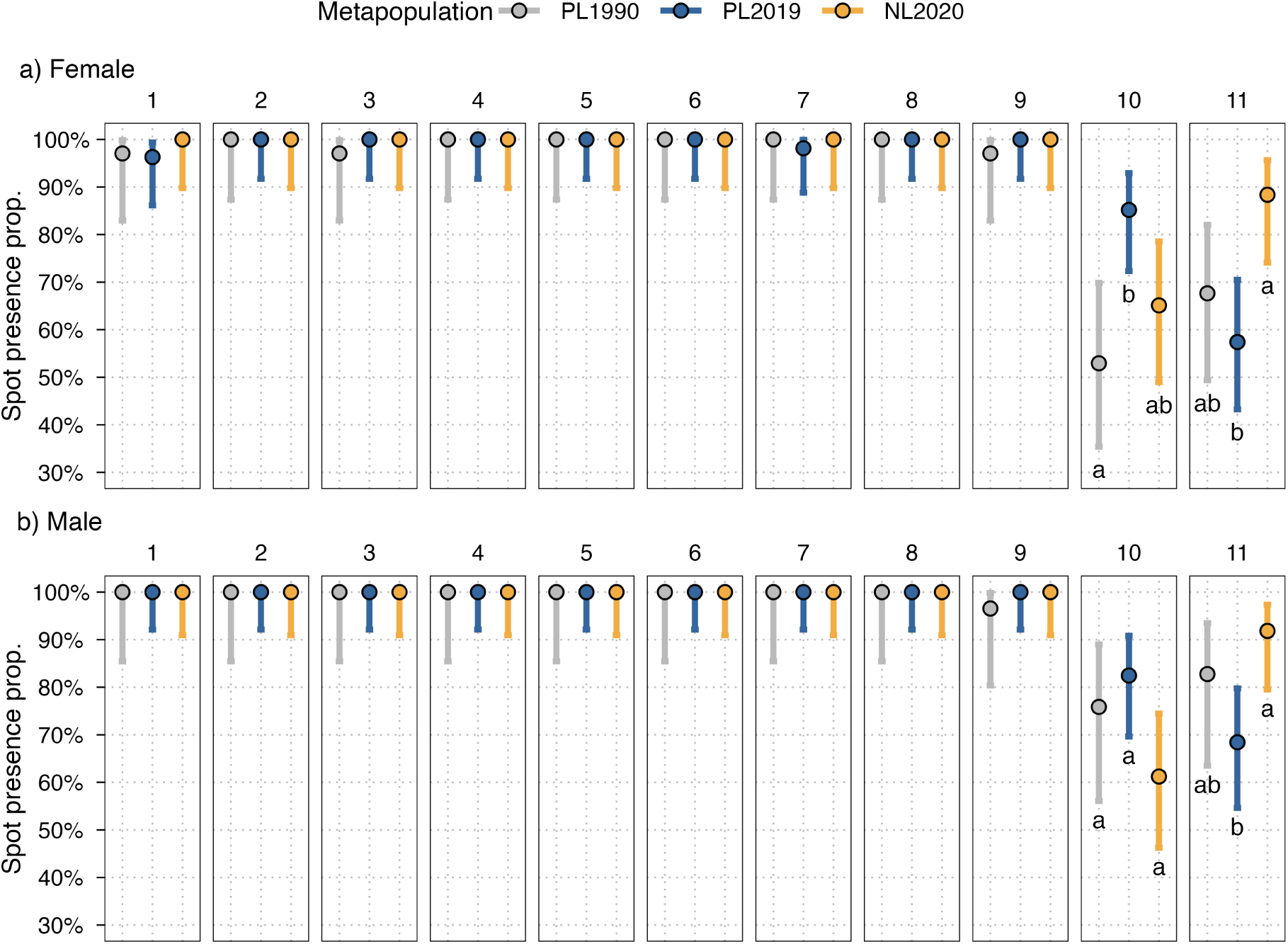
Proportion of spot presence in *P. teleius* a) females and b) males from the source (PL1990), current Polish (PL2019) and reintroduced Dutch (NL2020) metapopulations. Panels show the eleven wing spots; dots are proportions and bars their 95% confidence intervals (Wilson score intervals). Letters, shown only for the facultative spots 10 and 11 (the only ones modelled), indicate statistically significant differences between metapopulations.

Spot area increased with the spot pattern centroid size faster than isometry (a slope of 2) in females (allometric slope 2.75, 95% CI 2.17–3.33) and close to isometry in males (2.39, 95% CI 1.91–2.88). At equal size, the metapopulations did not differ in spot area overall (females: d.f. = 2, *χ*^2^ = 0.05, P = 0.975; males: d.f. = 2, *χ*^2^ = 0.40, P = 0.820), but they differed in particular spots (interaction, females: d.f. = 20, *χ*^2^ = 65.57, P < 0.001; males: d.f. = 20, *χ*^2^ = 101.18, P < 0.001; Fig. S4). Dutch butterflies had a larger spot 1 than those from the source metapopulation in both sexes (by 42% in females and 45% in males; and by 27% than the current Polish ones in males), and Dutch males had a smaller facultative spot 11 than source males (0.47 times as large). Dutch females also had a smaller spot 4 than source females, and Dutch males a smaller spot 10 than current Polish males. When the 17 (females) and 11 (males) observations with extreme residuals were left out, the differences in spots 1 and 11 remained, and the larger spot 1 of Dutch females also differed from the current Polish ones, whereas those in spots 4 and 10 did not (Table S1).

Total spot area also increased with size (slope 2.83, 95% CI 2.18–3.47, in females and 2.57, 95% CI 2.01–3.13, in males), and at equal size it did not differ among metapopulations (females: d.f. = 2, *χ*^2^ = 0.29, P = 0.864; males: d.f. = 2, *χ*^2^ = 0.00, P = 0.999; Fig. 4), also when tested with heteroscedasticity-robust errors (females P = 0.872; males P = 0.999).

**Fig. 4:**
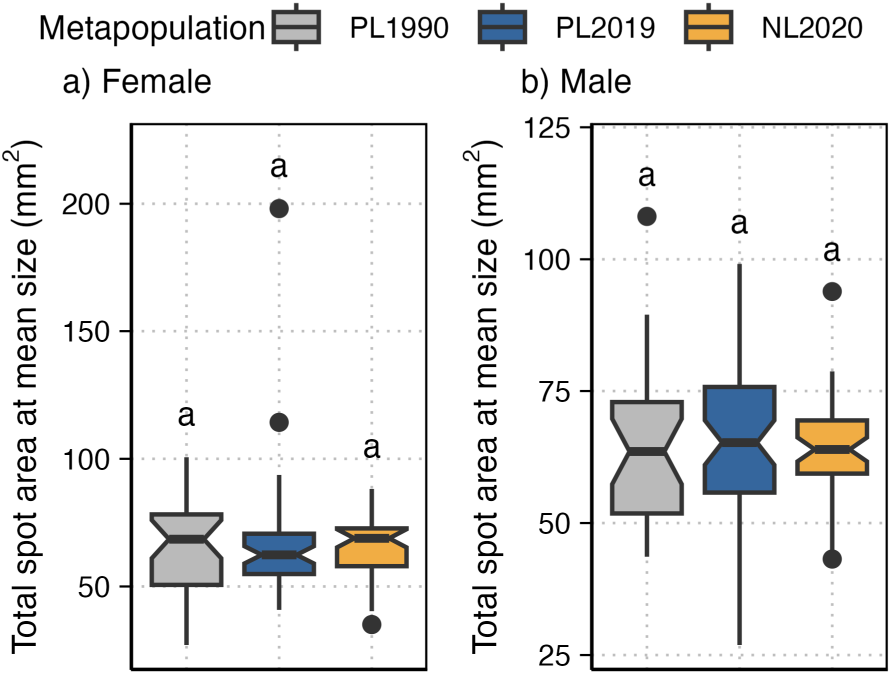
Total wing spot area standardized to the mean spot pattern centroid size of *P. teleius* a) females and b) males from the source (PL1990), current Polish (PL2019) and reintroduced Dutch (NL2020) metapopulations. Different letters at the top of the boxplots indicate statistically significant differences between groups. Boxes show the median and the interquartile range, whiskers extend to 1.5 times the interquartile range and notches show an approximate 95% confidence interval of the median; dots are values beyond the whiskers.

Spots 1 to 9 were symmetrical in nearly all butterflies (at least 90% of the females and 96% of the males of every metapopulation). In the facultative spots 10 and 11, the concordance between the presence of a spot on both wings differed among metapopulations and was lowest in the reintroduced Dutch metapopulation, in both sexes (left-right odds ratio, females: PL1990 52.7, PL2019 43.0, NL2020 3.2; males: PL1990 86.9, PL2019 23.4, NL2020 3.0; difference among metapopulations, females: d.f. = 2, *χ*^2^ = 7.55, P = 0.023; males: d.f. = 2, *χ*^2^ = 7.79, P = 0.020; Fig. 5). The pairwise differences were not significant after the Bonferroni correction (NL2020 against PL1990: females P = 0.094, males P = 0.071; against PL2019: females P = 0.112, males P = 0.151).

**Fig. 5:**
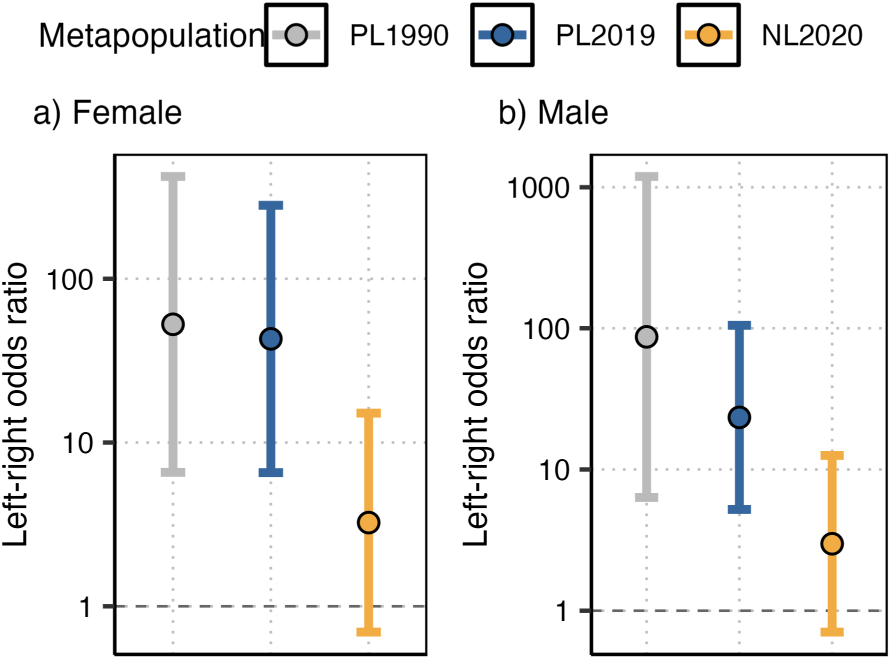
Concordance between the two hindwings in the presence of the facultative spots 10 and 11 of *P. teleius* a) females and b) males from the source (PL1990), current Polish (PL2019) and reintroduced Dutch (NL2020) metapopulations: odds ratio of the presence of a spot on the right wing given its presence on the left one (log scale), with its 95% confidence interval. High values indicate symmetrical butterflies; the dashed line (odds ratio of one) indicates no concordance between wings.

Among the individuals with both spots present, the proportion of spot fusion (between spots 8 and 11) in females did not differ significantly among metapopulations, being at the threshold of significance (metapopulation: d.f. = 2, *χ*^2^ = 5.98, P = 0.050; Fig. 6a). However, the proportion of spot fusion in males was significantly influenced by butterfly metapopulation (metapopulation: d.f. = 2, *χ*^2^ = 7.05, P = 0.029; Fig. 6b). Spot fusion was found in a significantly higher proportion in males from the Polish source metapopulation (PL1990) than in males from the current Polish metapopulation (PL2019). A similar but not statistically significant pattern was found for females.

**Fig. 6:**
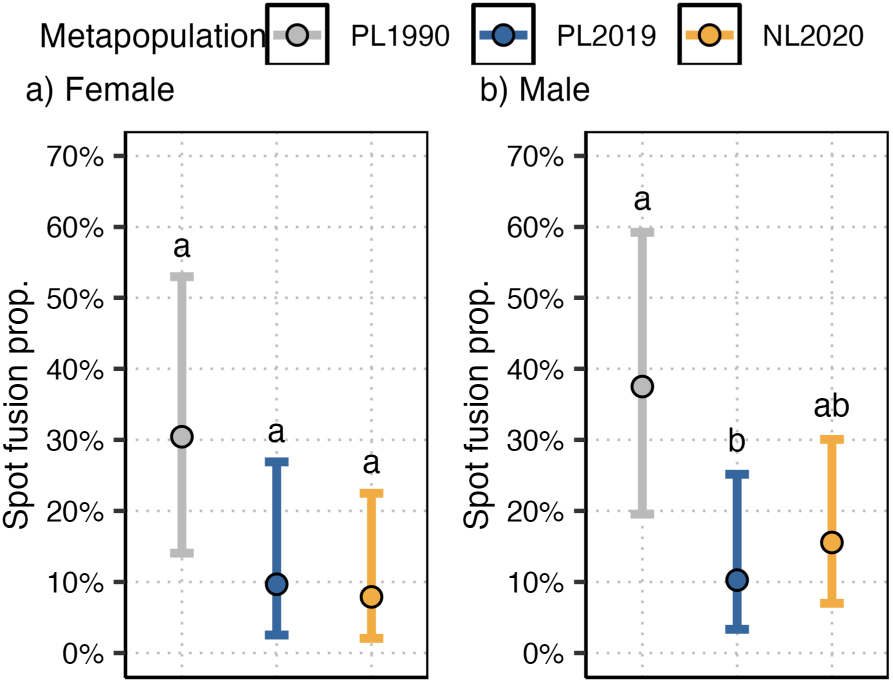
Proportion of wing spot fusion (spots 8 and 11) of *P. teleius* a) female and b) male butterflies with both spots present, from the source (PL1990), current Polish (PL2019) and reintroduced Dutch (NL2020) metapopulations. Bars are 95% confidence intervals (Wilson score intervals), and different letters indicate statistically significant differences between groups.

The PERMANOVA analysis showed a significant effect of the butterfly metapopulation on the wing spot pattern for both sexes (female, d.f. = 2; F = 2.97; P = 0.016; male, d.f. = 2; F = 3.47; P = 0.009; Fig. S5). The metapopulation also had a significant effect on the spot pattern variability, measured as the distance of each individual to the centroid of its metapopulation (female, d.f. = 2; F = 6.91; P = 0.001; male, d.f. = 2; F = 3.53; P = 0.031; Fig. 7), so the differences found by the PERMANOVA may partly reflect differences in the dispersion of the metapopulations and not only in their location. In both sexes, butterflies from the reintroduced Dutch metapopulation (NL2020) showed a significantly lower variability than those from the source metapopulation (PL1990; females P = 0.003; males P = 0.027), while the current Polish metapopulation (PL2019) was intermediate and did not differ significantly from either (females P = 0.087 and 0.390; males P = 0.840 and 0.249).

**Fig. 7:**
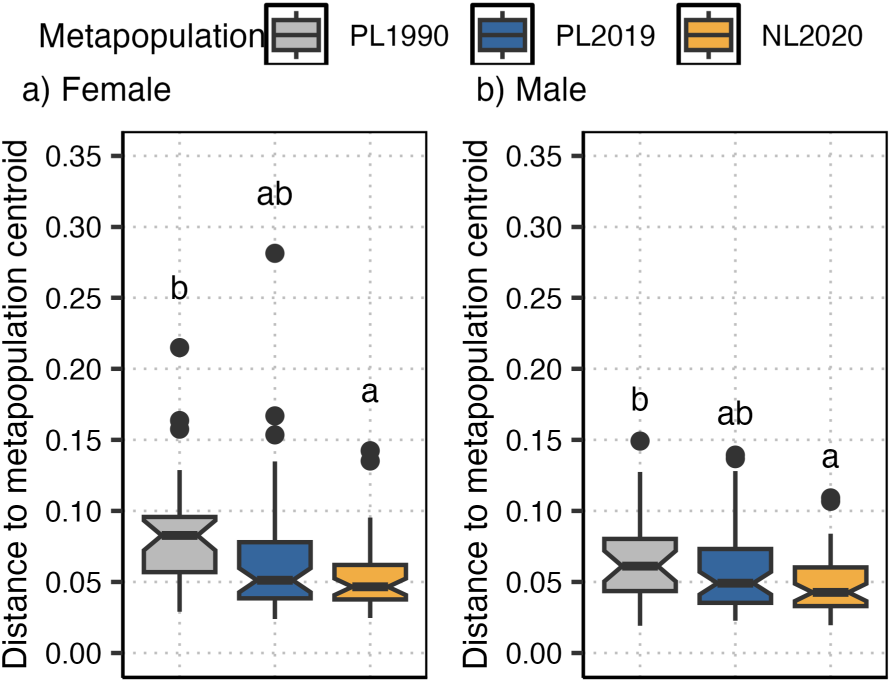
Spot pattern variability: Bray-Curtis distance of the wing spot pattern of each individual to the centroid of its metapopulation for a) females and b) males from the source (PL1990), current Polish (PL2019) and reintroduced Dutch (NL2020) metapopulations. Different letters indicate statistically significant differences between metapopulations. Boxes show the median and the interquartile range, whiskers extend to 1.5 times the interquartile range and notches show an approximate 95% confidence interval of the median; dots are values beyond the whiskers.

## Discussion

Our results revealed that the wing spot pattern in *P. teleius* butterflies can change over time in a specific metapopulation, but it can also differ after a reintroduction process. The spot pattern shape, and in males the size of spot 1, separated the reintroduced Dutch metapopulation from both Polish ones, as expected from the founder effect and genetic drift (hypothesis 1). The Dutch metapopulation’s lower spot pattern variability, the larger spot 1 of its females and the smaller spot 11 of its males differed only from the source metapopulation. Its more frequent spot 11 differed only from the current Polish one, and its lower concordance between wings in spot presence was significant only overall.

The spot pattern shape separated the reintroduced Dutch metapopulation from both Polish ones, which did not differ from each other, although in males the difference from the source metapopulation did not remain when only the nine permanent spots were used. Other studies have already reported differences in wing shape between metapopulations of the same butterfly species (Mikitová *et al*., 2022), and the hindwing shape of *P. teleius* changed in both current metapopulations (Sánchez-García *et al*., 2024), but the spot pattern shape differed only between the Dutch metapopulation and the Polish ones, so it did not simply follow the wing shape. Our results also revealed the highest spot pattern variability in the source Polish metapopulation from 1990 and the lowest one in the Dutch reintroduced metapopulation, with the current Polish metapopulation intermediate, although the Dutch metapopulation was only significantly less variable than the source one. Such results may reflect the loss of genetic variability. The Dutch metapopulation suffered a bottleneck and a consequent reduction of genetic richness due to the reintroduction, with half of the alleles of the current Polish metapopulation (Sánchez-García *et al*., 2024); this genetic comparison, however, was made with the current Polish metapopulation, from which the Dutch one did not differ significantly in spot pattern variability. Thus, despite the relatively high number of founders, only part of the variability of the source metapopulation may have been transferred to the Netherlands, and, as the Dutch metapopulation is isolated and has received no further butterflies since 1990, this variability could not be restored by gene flow. The Dutch butterflies also differed in particular spots, mostly from only one of the Polish samples. Spot 11 was more frequent in their wings than in those of the current Polish butterflies in both sexes, and, at equal size, they had a larger spot 1 than the source butterflies and, in males, also than the current Polish ones, whereas Dutch males had a smaller spot 11 than the source males (the smaller spot 4 of Dutch females and spot 10 of Dutch males did not remain without the extreme residuals). The differences from the current Polish metapopulation could be the effect of the genetic differentiation between the two current metapopulations (Sánchez- García *et al*., 2024). It is known that parent spot size is correlated with offspring spot presence in some species of butterflies, suggesting that the inheritance of the spot pattern could be described as a dosage model with different thresholds below which each spot is absent (Brakefield & van Noordwijk, 1985). Spot size is directly determined at the gene level (Holloway *et al*., 1993), but different spots from the same wing can be under different selective pressures (Obara & Rutowski, 2023), so these differences cannot be attributed to drift alone. They could also be due to adaptation or plastic responses to the local conditions of the Netherlands, with a cooler climate (Sánchez-García *et al*., 2024).

Differences in the presence of particular spots among populations have already been found in *Phengaris arion*, with no constant hindwing spot in Polish populations (Sielezniew & Dziekańska, 2011) and fewer and smaller spots in western Finnish populations (Väisänen *et al*., 1994). In our study, only the facultative spots 10 and 11 varied in presence. Only a few traits have changed in the current Polish metapopulation since 1990. Unlike spot 11, spot 10 changed in the source region. It was less frequent in females from the source metapopulation than in females from the current Polish metapopulation, while the Dutch females did not differ significantly from either. This change in the current Polish metapopulation, like its larger and differently shaped hindwings (Sánchez-García *et al*., 2024), could reflect local changes in the source region, where habitat patches have been lost and their connectivity has decreased under an increasing urbanization pressure (Kajzer-Bonk & Nowicki, 2022; Sánchez-García *et al*., 2024). The difference in the presence of spot 10 was found only in females, which agrees with the sex-dependent expression of particular spots in some butterflies (hindwing spots in *Maniola jurtina* (Brakefield & van Noordwijk, 1985); forewing spots in *Pieris rapae* (Stoehr *et al*., 2016)); moreover, in lycaenids the female hindwing spot pattern can work as a mate-recognition signal (Fordyce *et al*., 2002). Among the individuals with both spots present, the proportion of fusion between spots 8 and 11 was lower in both current metapopulations than in the source metapopulation from 1990, but the only significant difference was between the source and the current Polish metapopulation in males; as the Dutch metapopulation did not differ significantly from either, a change in the source region cannot be told apart from one shared by both regions. Short-term changes in wing spot size have been reported over a 23-year period in *Papilio polytes* (Katoh *et al*., 2017), but as the function of the spots in *P. teleius* is unknown, we cannot identify the factor behind these changes.

Total spot area did not differ among metapopulations once its allometric scaling with size was taken into account, so the larger butterflies of the current Polish metapopulation (Sánchez-García *et al*., 2024) did not have a more extensive spot pattern relative to their size. Furthermore, spot pattern shape changed with size (allometry), significantly in males and at the threshold of significance in females. Since these spots are on the ventral side of the hindwing, we did not expect them to play a role in thermoregulation, and our results give no evidence of a change in total melanization.

The wing spot pattern can also offer potential to study FA in relation to the environmental conditions. It has been proved that the eyespot symmetry can be sensitive to stress (Brakefield, 1997). Moreover, spot pattern FA does not generally show any indication of heritable variation; however, mutant individuals with additional spots seem to be more heritably influenced, exhibiting more non-directional asymmetry (Brakefield & Breuker, 1996). Here, the concordance between the two wings in the presence of the facultative spots differed among metapopulations, being lowest in the reintroduced Dutch metapopulation in both sexes and regardless of how often the spots were present, although no pairwise difference was significant after the Bonferroni correction. If confirmed, such higher FA could be explained by the increase of the stress level during the caterpillar development or by the loss of genetic variability due to the bottleneck that the reintroduced metapopulation suffered (Parsons, 1992; Sánchez-García *et al*., 2024), as expected from the founder effect. However, FA as an indicator of population conservation status should be carefully taken into account as changes in FA could also be a response to differences in environmental conditions (Windig *et al*., 2000). For instance, neither significant rise in asymmetry nor decline of morphological variability were detected after several bottlenecks in a population of *Parnassius apollo* (Habel *et al*., 2012).

The butterflies from the source metapopulation from 1990 were those that died during the trip to the Netherlands, and they could therefore be biased towards weaker individuals (Sánchez-García *et al*., 2024). However, several observations argue against such a bias. Their wings were smaller than those from the current Polish metapopulation, but did not differ in size from those of butterflies collected at random in 2003 and 1996 (Sánchez-García *et al*., 2024), and size was a covariate in the analyses of spot pattern shape and spot area. Moreover, the spot pattern shape of the source metapopulation did not differ significantly from that of the current Polish metapopulation, which would be unlikely if butterflies with particular spot patterns had died more often during the trip. Wing size was not related to survival in experiments with the butterfly *Pontia occidentalis* either (Kingsolver, 1999). A bias of this sample cannot be ruled out entirely; it would affect the comparisons with the source metapopulation (the difference in spot pattern shape between the Dutch and source metapopulations, the lower spot pattern variability of the Dutch metapopulation, the larger spot 1 of Dutch females, the smaller spot 11 of Dutch males and the changes in spot 10 and in spot fusion), but not those between the two current metapopulations.

Our results have some implications for the conservation of reintroduced insect populations, although they come from a single reintroduction, without replication. The reintroduction of 86 butterflies was enough for *P. teleius* to persist and expand in the Netherlands for about thirty generations (Sánchez-García *et al*., 2024), but its wing spot pattern kept only part of the variability of the source metapopulation, and its genetic variation only part of that of the current Polish one (Sánchez-García *et al*., 2024). As the Dutch metapopulation is isolated, this variability cannot be restored by gene flow. The spot pattern variability, which can be measured non-invasively from photographs of the hindwing underside, could complement the genetic monitoring of reintroduced populations if it is confirmed to track genetic variability, whereas FA should be used with caution as an indicator of their status (Habel *et al*., 2012; Windig *et al*., 2000). Moreover, as the source metapopulation also changed after the reintroduction, following both the source and the reintroduced populations, and keeping a reference sample of the translocated individuals, are needed to separate the effects of the reintroduction from changes over time.

To conclude, the wing spot pattern of *P. teleius* changed in the thirty years after the reintroduction, although not every trait differed in both sexes or in every pairwise comparison. The spot pattern shape separated the reintroduced Dutch metapopulation from both Polish ones, whereas most of its other differences, including its lower spot pattern variability, were with only one of the Polish metapopulations or only overall, which agrees with the founder effect and the loss of genetic variability after the reintroduction (Sánchez-García *et al*., 2024). Only the presence of spot 10 in females and the spot fusion in males changed in the current Polish metapopulation, which could reflect environmental changes over time in the source region, although for spot fusion a change shared by both regions cannot be ruled out. As adaptation or plastic responses to the local conditions of the Netherlands could produce differences similar to those of drift, rearing butterflies from both metapopulations in a common environment, or comparing the differentiation of the spot pattern with that of neutral genetic markers (QST–FST comparisons, Leinonen *et al*., 2013), would be needed to separate them.

## Acknowledgements

The study was funded by the Polish National Science Centre (NCN) grant 2018/31/B/NZ8/03476. IW received additional funds from the Province of Northern Brabant, the Netherlands. National State Forestry, Natuurmonumenten and the Province of Northern Brabant gave us permission to access the nature reserve and carry out the survey. Permission for butterfly capture in Kraków was given by the Regional Directorate for Environmental Protection in Kraków (decision OP-I.6401.156.2019.KW). VC was supported by the Polish National Science Centre (NCN) grant 2022/47/D/NZ8/01758. We would like to thank Istvan Maak for his help during the fieldwork. We would also like to thank Cristina Sevilleja and Juan Gallego Zamorano for their help during the fieldwork in Moerputten.

## Author contributions

**Daniel Sánchez-García:** Conceptualization; methodology; investigation; data curation; formal analysis; software; validation; visualization; writing – original draft; writing – review and editing. **Luca Pietro Casacci:** Conceptualization; methodology; investigation; writing – review and editing; supervision. **Irma Wynhoff:** Conceptualization; methodology; investigation; resources; writing – review and editing. **Violette Chiara:** Methodology; software; writing – review and editing. **Magdalena Witek:** Conceptualization; methodology; investigation; resources; writing – review and editing; supervision; project administration; funding acquisition.

## Data availability statement

The raw data and the R code that reproduces the analyses are available in Zenodo at https://doi.org/10.5281/zenodo.23959599.

