## Supporting information for "Changes in the wing spot pattern of the endangered butterfly *Phengaris teleius* thirty years after its reintroduction"

### Appendix S1. Computer assisted spot detection

We developed a program in Python (3.9) to measure the area and the exact centre coordinates of the black spots from the hindwing images. The image management and treatment were based on the opencv library (Bradski, 2000). All images were first rotated and resized so that the scale and orientation were the same for all pictures. Images were converted to grayscale and blurred to prevent the wing scales from biasing the detection. The lighting (average value) of the images was corrected so that each butterfly wing would have the same average level of brightness. Black spots were identified from these images using an adaptive thresholding by applying the adaptiveThreshold() function from the opencv package (Bradski, 2000). Those spots were also filtered by size, to discard objects smaller or larger than the wing spots, such as noise or shadows. Finally, we associated these automatically detected spots with the manually recorded spot positions using the Hungarian algorithm provided by the linear_sum_assignment() function from the scipy package (Virtanen *et al.*, 2020). We also determined whether spots were detected in the wing areas in which facultative spots were expected (i.e. spots 10 and 11).

A Graphical User Interface was also programmed to allow manual corrections of the errors made by the automatic detection program. At this stage, users could redraw the areas of the detected spots, remove or add facultative spots, or reassign them. After this correction, the area and centroid (arithmetic mean) of each of the spots were calculated. Spots 8 and 11 were considered fused when the program detected them as two separate spots joined to each other.

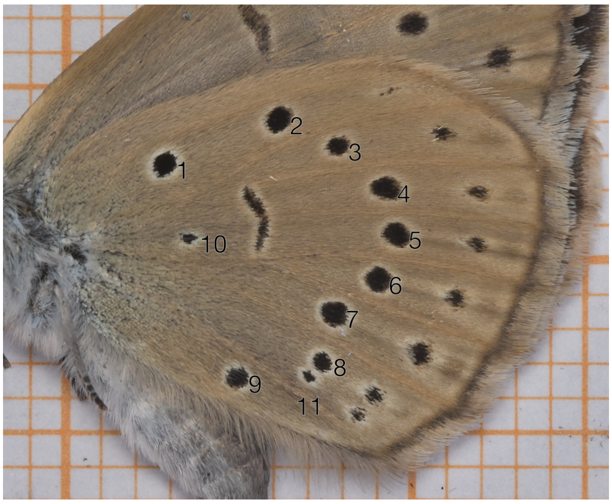


Fig. S1: Hindwing underside of a female *P. teleius* from the reintroduced Dutch metapopulation (NL2020), with the spots numbered as in Fig. 1. Spots 1–9 are permanent and spots 10 and 11 facultative. The background is millimetre paper.

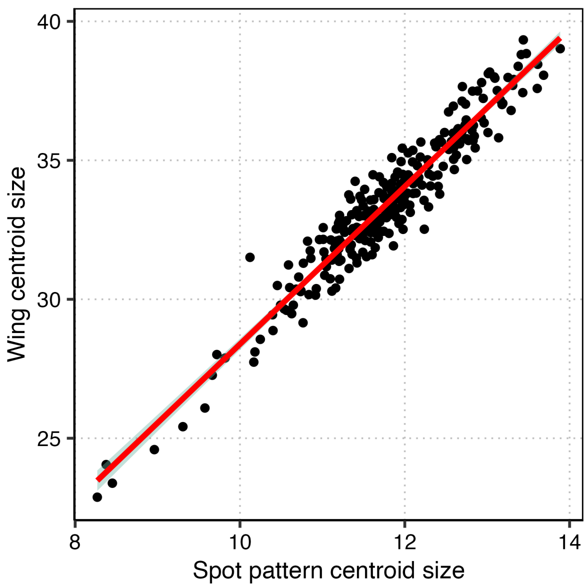


Fig. S2: Correlation plot of spot pattern centroid size and wing centroid size of the hindwings, for females and males pooled (n = 266 butterflies).

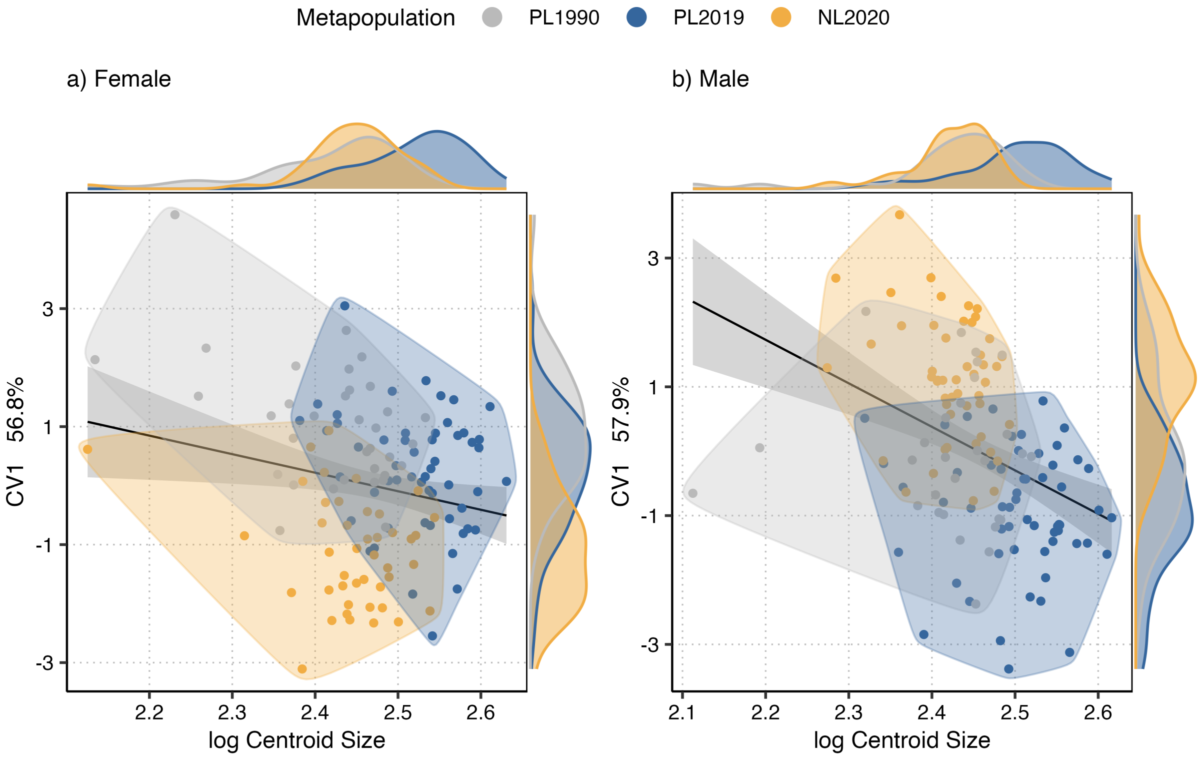


Fig. S3: Spot pattern shape allometry of *P. teleius* a) female and b) male butterflies from the Polish and Dutch metapopulations in different years: PL1990 (grey: year of the reintroduction), PL2019 (blue: current Polish metapopulation) and NL2020 (yellow: reintroduced Dutch metapopulation). First canonical variate (CV1) of the spot pattern shape against the logarithm of the spot pattern centroid size; the line, with its 95% confidence band, is a single regression for all metapopulations, shaded areas enclose the individuals of each metapopulation, and the axis density plots show the distribution of each variable.

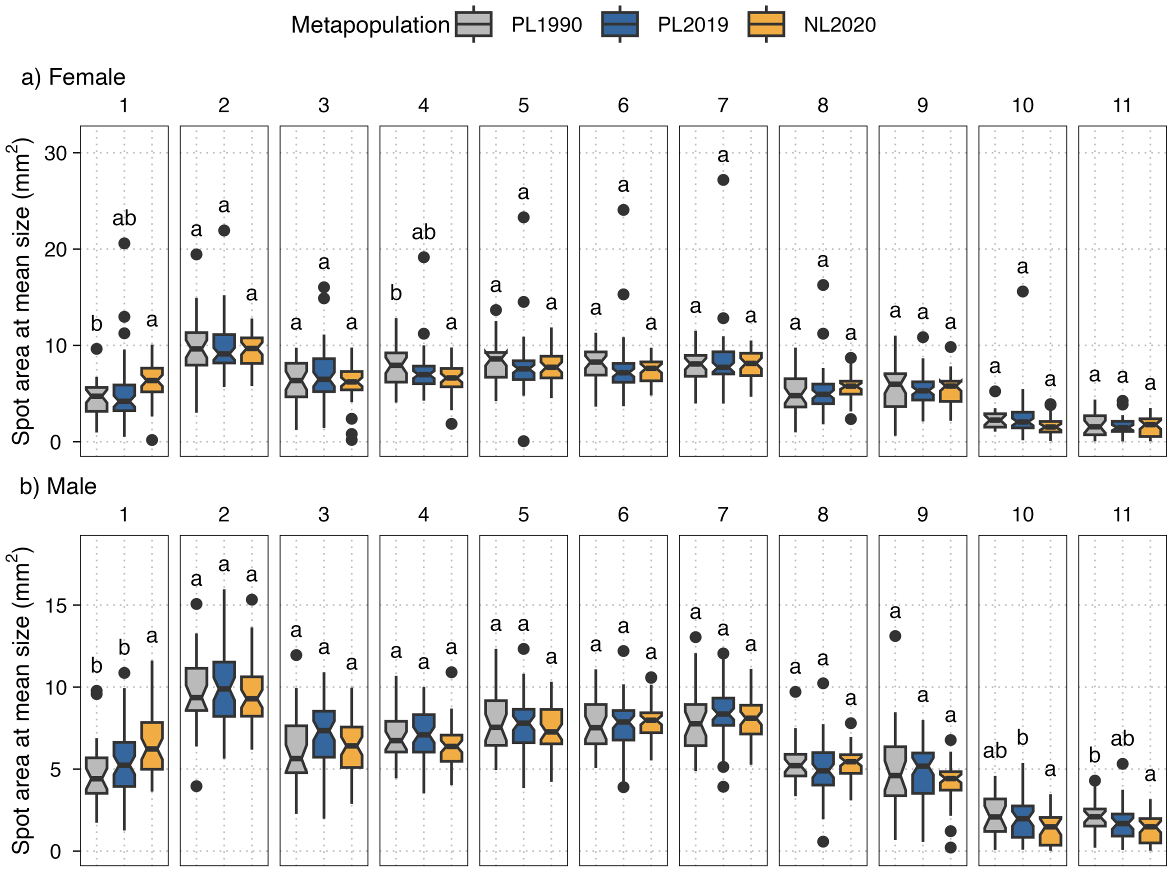


Fig. S4: Area of each wing spot standardized to the mean spot pattern centroid size of *P. teleius* a) females and b) males from the source (PL1990), current Polish (PL2019) and reintroduced Dutch (NL2020) metapopulations. Panels show the eleven wing spots and boxes the metapopulations. Different letters at the top of the boxplots indicate statistically significant differences between metapopulations. Boxes show the median and the interquartile range, whiskers extend to 1.5 times the interquartile range and notches show an approximate 95% confidence interval of the median; dots are values beyond the whiskers.

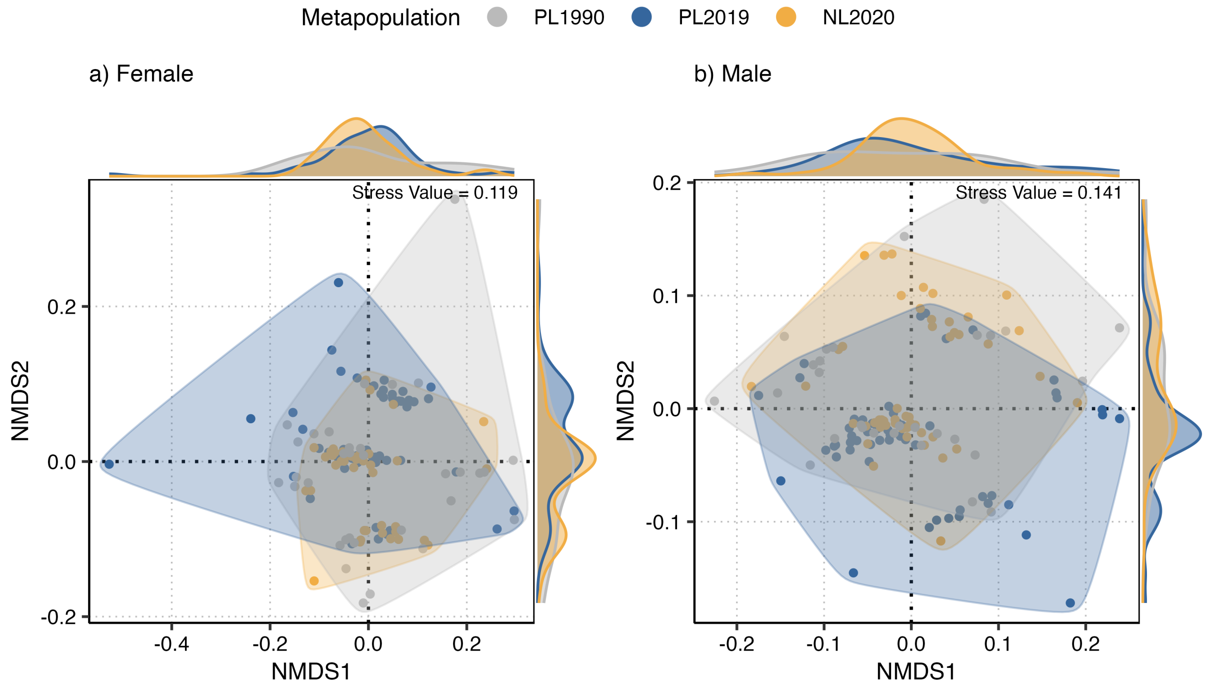


Fig. S5: Non-metric multidimensional scaling (NMDS) representation of the Bray-Curtis distances, based on spot presence, size-standardized spot area and fusion, of *P. teleius* a) female and b) male butterfly wing spot pattern coming from the Polish source (PL1990), current Polish (PL2019) and Dutch (NL2020) metapopulations. Shaded areas enclose the individuals of each metapopulation, the axis density plots represent the distribution of each NMDS component, and the stress value of each ordination is given in its panel.

Table S1: Assumption checks of the models of spot presence, asymmetry, fusion, spot area and total spot area, of spot pattern shape and of the spot pattern dissimilarity, for females and males, including the spot area model refitted without its extreme residuals. Binomial models: simulated residuals with the R package DHARMa (1000 simulations; uniformity, dispersion and outlier tests). Gaussian models: distribution of the residuals (Shapiro-Wilk test for total spot area; number of normalised residuals beyond |3| and excess kurtosis for the spot area model) and Levene's test, centred on the median, of their variance among metapopulations; total spot area also with a Wald test with heteroscedasticity-robust (HC3) standard errors. Procrustes ANOVAs, which assume no distribution: homogeneity of shape disparity (Procrustes variance) among metapopulations.

| Analysis | Sex | Check |
|---|---|---|
| Spot presence (binomial GLMM) | Females | Simulated residuals (DHARMa): uniformity P = 0.117, dispersion P = 0.748, outliers P = 1.000 |
| | Males | Simulated residuals (DHARMa): uniformity P = 0.731, dispersion P = 0.154, outliers P = 1.000 |
| Spot asymmetry, left-right concordance (binomial GLMM) | Females | Simulated residuals (DHARMa): uniformity P = 0.356, dispersion P = 0.110, outliers P = 1.000 |
| | Males | Simulated residuals (DHARMa): uniformity P = 0.444, dispersion P = 0.196, outliers P = 1.000 |
| Spot fusion (binomial GLM) | Females | Simulated residuals (DHARMa): uniformity P = 0.367, dispersion P = 0.928, outliers P = 1.000 |
| | Males | Simulated residuals (DHARMa): uniformity P = 0.898, dispersion P = 0.916, outliers P = 1.000 |
| Spot area (LMM) | Females | Normalised residuals: 17 of 1357 beyond \|3\| (excess kurtosis 18.3); equal variance among metapopulations (Levene) P = 0.492 (one residual variance per spot is estimated); refitted without the extreme residuals, metapopulation P = 0.892, interaction P < 0.001, significant pairwise contrasts: spot 1, NL2020 - PL1990 (ratio 1.53); spot 1, NL2020 - PL2019 (ratio 1.30) |
| | Males | Normalised residuals: 11 of 1420 beyond \|3\| (excess kurtosis 4.4); equal variance among metapopulations (Levene) P = 0.329 (one residual variance per spot is estimated); refitted without the extreme residuals, metapopulation P = 0.809, interaction P < 0.001, significant pairwise contrasts: spot 1, NL2020 - PL1990 (ratio 1.45); spot 1, NL2020 - PL2019 (ratio 1.24); spot 11, NL2020 - PL1990 (ratio 0.51) |
| Total spot area (Gaussian GLM) | Females | Residual normality (Shapiro-Wilk) P = 0.001; equal variance among metapopulations (Levene) P = 0.046; metapopulation with heteroscedasticity-robust errors (HC3) P = 0.872 |
| | Males | Residual normality (Shapiro-Wilk) P = 0.011; equal variance among metapopulations (Levene) P = 0.010; metapopulation with heteroscedasticity-robust errors (HC3) P = 0.999 |
| Spot pattern shape (Procrustes ANOVA) | Females | Residual randomization (RRPP), no distributional assumption; equal shape disparity (Procrustes variance) among metapopulations, Bonferroni-adjusted pairwise P: PL1990–NL2020 0.510; PL2019–NL2020 1.000; PL2019–PL1990 1.000 |
| | Males | Residual randomization (RRPP), no distributional assumption; equal shape disparity (Procrustes variance) among metapopulations, Bonferroni-adjusted pairwise P: PL1990–NL2020 1.000; PL2019–NL2020 1.000; PL2019–PL1990 1.000 |
| Spot pattern [illegible] | Females | Homogeneity of multivariate dispersion (PERMDISP) P = 0.001: dispersion differs [illegible] |

Table S2: Estimated marginal means (EMMs) pairwise comparisons of the female and male wing spot pattern shape between the source (PL1990), current Polish (PL2019) and reintroduced Dutch (NL2020) metapopulations (Bonferroni-adjusted P values).

| | | PL1990 | PL2019 |
|---|---|---|---|
| Females | NL2020 | Z = 2.94<br>P = 0.006** | Z = 3.14<br>P = 0.003** |
| | PL1990 | - | Z = 1.39<br>P = 0.249 |
| Males | NL2020 | Z = 2.25<br>P = 0.045* | Z = 3.49<br>P = 0.003** |
| | PL1990 | - | Z = 1.93<br>P = 0.084 |
